# Large-scale wearable data reveal spatiotemporal organization of annual sleep patterns

**DOI:** 10.1101/2025.08.03.668323

**Authors:** Annika H. Rose, Hannes Schenk, Benjamin F. Maier, Dirk Brockmann, Eva C. Winnebeck

**Author notes:** Corresponding author(s). E-mail(s): annika. Contributing authors.

## Abstract

Sleep is fundamental to health, yet large-scale, objective data on how geography shapes sleep behavior remain scarce. We analyzed over 45 million nights of sensor data from 105,741 German adults wearing consumer-grade wearables across 2.7 years. Sleep timing displayed a continuous east–west gradient, with later onset, midsleep, and offset in western regions, consistent with solar progression. This effect was strongest on weekends and in rural areas, where midsleep was delayed by 2.2 minutes per degree longitude and sleep duration increased by 1.0 minute. A north–south gradient also emerged. Weekday midsleep advanced by 0.9 minutes per degree latitude, while weekend midsleep was delayed by 0.2 minutes, resulting in greater social jetlag in the north. Sleep duration declined toward higher latitudes across both day types.

Seasonal analyses revealed consistent annual rhythms. Sleep duration increased by 24.7 minutes in winter relative to summer, and sleep offset closely followed sunrise. These patterns highlight the joint influence of solar and social time on sleep, with implications for regionally tailored public health strategies.

## 1 Introduction

Sleep is increasingly recognized as a cornerstone of human health, alongside physical activity, nutrition, and mental well-being. Far from a passive state, sleep plays an active and essential role in regulating immune, metabolic, cognitive, and emotional processes. Insufficient or disrupted sleep has been linked to a wide range of adverse outcomes, including cardiovascular and metabolic disease, psychiatric disorders, and premature mortality, and may act as a direct causal factor rather than merely a secondary symptom in many of these conditions [1, 2]. Moreover, sleep represents a modifiable risk factor and thus offers an important target for interventions aimed at preventing disease and enhancing public health [3–6].

Despite growing scientific consensus, sleep health remains undervalued in public health policy and under strain in modern 24-hour societies. The pressure to prioritize productivity over rest contributes to widespread sleep loss, often normalized despite its considerable health risks [7, 8]. Globally, around 35% of adults do not meet recommended sleep durations,, with marginalized populations disproportionately affected [9, 10]. Only 43 of 194 WHO member states have published population-level sleep data [1], reflecting a critical gap in surveillance and policy guidance. The societal costs are substantial: insufficient sleep is estimated to cost the United States $680 billion and Germany $55 billion annually through lost productivity, healthcare expenses, and safety risks [11]. Sleep deprivation has also been implicated in high-profile industrial disasters, such as the Chernobyl explosion and the Challenger accident, underscoring its broad relevance beyond individual health.

In response, sleep research has expanded beyond clinical disorders towards a broader, multidimensional conceptualization of sleep health, encompassing timing, quality, and efficiency. This shift reflects recognition of sleep as shaped by a complex interplay of behavioral, demographic, environmental, and psychosocial factors. Lower socioeconomic status, light and noise pollution, and chronic stress disproportionately undermine sleep health among disadvantaged populations [12, 13]. Understanding how these factors interact across time and space is essential for designing equitable public health interventions.

Recent advances in wearable technology have transformed sleep research by enabling high-resolution, longitudinal monitoring of sleep behavior across large populations. Unlike traditional methods that rely on self-reports or limited cross-sectional samples, wearable devices collect continuous, objective data over extended periods [14–16]. Prior research has shown that sleep timing tends to align with solar time, even in highly industrialized modern societies, suggesting persistent circadian entrainment to natural light cycles [17–20]. However, most of this evidence comes from questionnaire-based or diary studies with limited temporal and spatial resolution. Wearable-derived data now make it possible to revisit these patterns with greater precision and scale, allowing systematic assessment of how environmental and social timing jointly influence sleep behavior.

In this study, we analyze over 45 million nights of sleep data from more than 100,000 German residents who participated in the Corona Data Donation Project. Using consumer wearable data collected between April 2020 and December 2022, we quantify how sleep timing and duration vary across geography and season. This dataset represents the largest known objective record of sleep behavior in Germany and provides an unprecedented opportunity to examine population-level circadian patterns in situ.

Our results reveal consistent east–west differences in sleep timing, with later sleep observed in western regions despite a shared time zone, consistent with solar gradients. We also identify a novel north–south pattern, with greater weekday–weekend variation in the north, indicative of heightened social jetlag. Seasonal oscillations show longer, later sleep in winter and shorter, earlier sleep in summer, suggesting entrainment to photoperiod. Together, these findings illustrate the combined influence of solar and social time on sleep behavior.

By systematically mapping spatiotemporal sleep patterns, our study highlights how solar cues and social structures interact to shape circadian rhythms. These insights can inform targeted interventions such as adjusting school start times, promoting flexible work schedules, or mitigating artificial light exposure, with the goal of reducing circadian misalignment and promoting sleep health at the population level.

## 2 Results

We analyzed spatial and temporal variation in two core dimensions of sleep: *sleep timing* and *sleep duration*. Sleep timing was assessed using midsleep, defined as the the midpoint between sleep onset and offset, as a primary marker of circadian phase. We focused on weekend nights, where midsleep is less constrained by work and school schedules and better reflects individuals’ endogenous circadian preference. Sleep duration was calculated as the interval from sleep onset to offset, excluding interim wake periods. Sleep onset and offset times were also examined to provide additional context. Distributions of all sleep variables across weekdays and weekends are shown in Figure S1, confirming normality and robust differences by day type. To capture these systematic differences, all analyses were performed separately for weekends and weekdays.

The analytic sample included 105,741 participants who met the inclusion criteria (detailed in the Supplementary Methods). Collectively, they contributed over 46 million nights of wearable-derived sleep data. Descriptive statistics on age, sex, BMI, and sleep measures are summarized in Table 1, with regional demographics shown in Figure S2.

**Table 1.** Summary statistics for participant demographics and sleep variables in the demographic (N = 59,774) and maximal (N = 105,741) cohorts. Values are reported as mean (standard deviation) and percentage of non-missing observations (*N*, %). Sleep onset, midsleep, and offset are shown in HH:MM format; negative onset times indicate times before midnight. Percentages are calculated relative to the total number of participants in each cohort.

| Variable | Demographic Cohort |  | Maximal Cohort |  |
| --- | --- | --- | --- | --- |
|  | Mean (SD) | N (%) | Mean (SD) | N (%) |
| **Demographics** |  |  |  |  |
| Age (yrs) | 49.19 (12.75) | 59774 (100.0) | 49.15 (12.75) | 60903 (57.6) |
| BMI (kg/m<sup>2</sup>) | 26.16 (4.76) | 59774 (100.0) | 26.16 (4.77) | 60053 (56.8) |
| Male (N, %) | 30335 (50.7%) | - | 30611 (50.6%) | - |
| Female (N, %) | 29439 (49.3%) | - | 29880 (49.4%) | - |
| **Weekdays** |  |  |  |  |
| Duration (min) | 422.26 (71.59) | 58252 (97.3) | 425.77 (72.20) | 102794 (97.0) |
| Onset (hh:mm) | 23:09 (01:07) | 57457 (96.2) | 23:08 (01:07) | 101084 (96.3) |
| Offset (hh:mm) | 06:44 (01:14) | 57866 (96.7) | 06:44 (01:14) | 102184 (96.7) |
| Midsleep (hh:mm) | 02:56 (01:04) | 57870 (96.7) | 02:56 (01:04) | 102239 (96.8) |
| **Weekends** |  |  |  |  |
| Duration (min) | 454.44 (81.68) | 58258 (97.5) | 457.31 (82.49) | 102998 (97.3) |
| Onset (hh:mm) | 23:38 (01:23) | 57744 (96.6) | 23:37 (01:23) | 102126 (96.6) |
| Offset (hh:mm) | 07:54 (01:19) | 57669 (96.3) | 07:53 (01:19) | 102165 (96.4) |
| Midsleep (hh:mm) | 03:47 (01:11) | 57874 (96.7) | 03:47 (01:11) | 102234 (96.8) |

**Table 2.** Results from linear mixed-effects models predicting sleep timing and duration on weekends. Separate models were fit for each outcome variable, with participant-level random intercepts to account for repeated measures. Models included demographic, geographic, seasonal, and environmental predictors. All continuous variables were mean-centered prior to fitting. Coefficients are unstandardized and reported in minutes, with 95% confidence intervals in parentheses. Positive estimates reflect later sleep or longer duration; negative estimates indicate earlier sleep or shorter duration. For longitude, negative values represent westward delays; for latitude, positive values indicate earlier sleep or shorter durations in more northern districts. “Observations” reflects the number of daily records; “Num. Users” denotes the number of unique participants. Model fit is reported using AIC, BIC, and log-likelihood. Marginal and conditional *R*^2^ values indicate variance explained by fixed effects alone and by the full model, respectively, following Nakagawa et al. [23]. P-values were calculated using the Satterthwaite approximation. Thresholds and asterisk notation are defined below the table.

|  | *Sleep Variable* |  |  |  |
| --- | --- | --- | --- | --- |
|  | ONSET | MIDSLEEP | OFFSET | DURATION |
| Intercept | -40.680***<br>(-41.460, -39.840) | 209.580***<br>(208.860, 210.300) | 460.860***<br>(460.080, 461.640) | 456.946***<br>(456.263, 457.629) |
| Age | -0.420***<br>(-0.480, -0.420) | -0.600***<br>(-0.660, -0.600) | -0.840***<br>(-0.840, -0.780) | -0.415***<br>(-0.443, -0.387) |
| Sex[male] | 16.620***<br>(15.720, 17.460) | 8.880***<br>(8.100, 9.660) | 1.140**<br>(0.300, 2.040) | -4.971***<br>(-5.696, -4.247) |
| BMI | 0.660***<br>(0.540, 0.780) | 0.300***<br>(0.180, 0.360) | -0.060<br>(-0.120, 0.060) | -1.009***<br>(-1.083, -0.935) |
| Urbanization[urban] | 11.940***<br>(10.920, 12.900) | 11.280***<br>(10.380, 12.180) | 10.620***<br>(9.660, 11.640) | -0.973*<br>(-1.818, -0.127) |
| DST[yes] | 8.040***<br>(7.800, 8.280) | 8.340***<br>(8.160, 8.520) | 8.760***<br>(8.580, 9.000) | 0.108<br>(-0.123, 0.339) |
| Season[spring] | 3.000***<br>(2.760, 3.180) | 4.560***<br>(4.380, 4.740) | 6.360***<br>(6.120, 6.540) | 4.512***<br>(4.280, 4.745) |
| Season[summer] | -2.640***<br>(-3.240, -2.040) | -3.300***<br>(-3.780, -2.820) | -3.900***<br>(-4.440, -3.360) | -1.508***<br>(-2.130, -0.887) |
| Season[winter] | 1.440***<br>(1.020, 1.860) | 2.160***<br>(1.800, 2.520) | 3.180***<br>(2.760, 3.540) | 1.530***<br>(1.088, 1.972) |
| Longitude | -2.460***<br>(-2.760, -2.160) | -2.160***<br>(-2.400, -1.860) | -1.800***<br>(-2.100, -1.560) | 1.011***<br>(0.773, 1.250) |
| Latitude | 0.180<br>(-0.060, 0.420) | 0.240*<br>(0.000, 0.480) | 0.360**<br>(0.120, 0.600) | -0.570***<br>(-0.781, -0.359) |
| Sunrise | -4.860***<br>(-4.980, -4.740) | -0.540***<br>(-0.660, -0.420) | 3.780***<br>(3.660, 3.900) | 7.968***<br>(7.817, 8.119) |
| Spring:Lon | 0.840***<br>(0.770, 0.910) | 0.540***<br>(0.490, 0.590) | 0.300***<br>(0.240, 0.360) | -0.473***<br>(-0.546, -0.400) |
| Summer:Lon | -0.180***<br>(-0.260, -0.100) | -0.360***<br>(-0.420, -0.300) | -0.540***<br>(-0.600, -0.480) | -0.401***<br>(-0.470, -0.332) |
| Winter:Lon | 1.260***<br>(1.190, 1.330) | 1.080***<br>(1.020, 1.140) | 0.960***<br>(0.890, 1.030) | -0.184***<br>(-0.260, -0.108) |
| Lon:Urban | 0.960***<br>(0.480, 1.430) | 0.900***<br>(0.540, 1.260) | 0.900***<br>(0.420, 1.380) | -0.244***<br>(-0.599, 0.111) |
| Spring:Sunrise | 9.300***<br>(9.060, 9.540) | 6.600***<br>(6.360, 6.840) | 4.080***<br>(3.840, 4.320) | -5.386***<br>(-5.647, -5.125) |
| Summer:Sunrise | -0.840***<br>(-1.200, -0.480) | -2.880***<br>(-3.240, -2.520) | -4.860***<br>(-5.210, -4.510) | -4.778***<br>(-5.162, -4.394) |
| Winter:Sunrise | 13.800***<br>(13.440, 14.160) | 12.120***<br>(11.760, 12.480) | 10.500***<br>(10.150, 10.850) | -2.259***<br>(-2.651, -1.867) |
| *Observations* | 7,306,659 | 7,315,838 | 7,284,966 | 7,369,356 |
| *Num. Users* | 59,772 | 59,772 | 59,772 | 59,772 |
| *Log Likelihood* | -11,061,576 | -9,545,282 | -10,287,102 | -41,718,914 |
| *AIC* | 22,123,194 | 19,090,605 | 20,574,245 | 83,437,870 |
| *BIC* | 22,123,484 | 19,090,895 | 20,574,535 | 83,438,160 |
| *Marg./Cond. R<sup>2</sup>* | 0.03/0.48 | 0.02/0.50 | 0.01/0.45 | 0.01/0.33 |

Note:
\* $p < 0.05$ ; \*\* $p < 0.01$ ; \*\*\* $p < 0.001$ .

Age and BMI were both associated with sleep behavior (Fig. S3). Older individuals exhibited earlier sleep onset, midsleep, and offset times, along with shorter sleep durations. Similarly, higher BMI was linked to slightly delayed sleep timing and modest reductions in duration. These effects were broadly consistent across sexes and day types, with greater variability on weekends. These patterns align with prior research on age- and weight-related differences in sleep architecture and circadian phase [21, 22].

To ensure robustness, we performed sensitivity analyses across demographic subgroups and alternative model specifications (see Supplementary Information). The findings remained stable under varied assumptions, supporting the reliability of the main results.

### 2.1 Longitudinal gradients in sleep — Earlier and longer sleep in the east compared to the west

Visual inspection of district-aggregated sleep data (Fig. 1A-B) revealed a clear east-west gradient in sleep timing, with individuals in eastern Germany sleeping earlier than those in western regions across all timing measures. Several large metropolitan areas stood out with consistently later sleep than surrounding districts, forming localized hotspots (Fig. S4). Visualizations were stratified by urbanization to assess urban–rural differences, and by day type to account for weekly variation in sleep pattern (Fig. 1B–F).

**Fig. 1.**
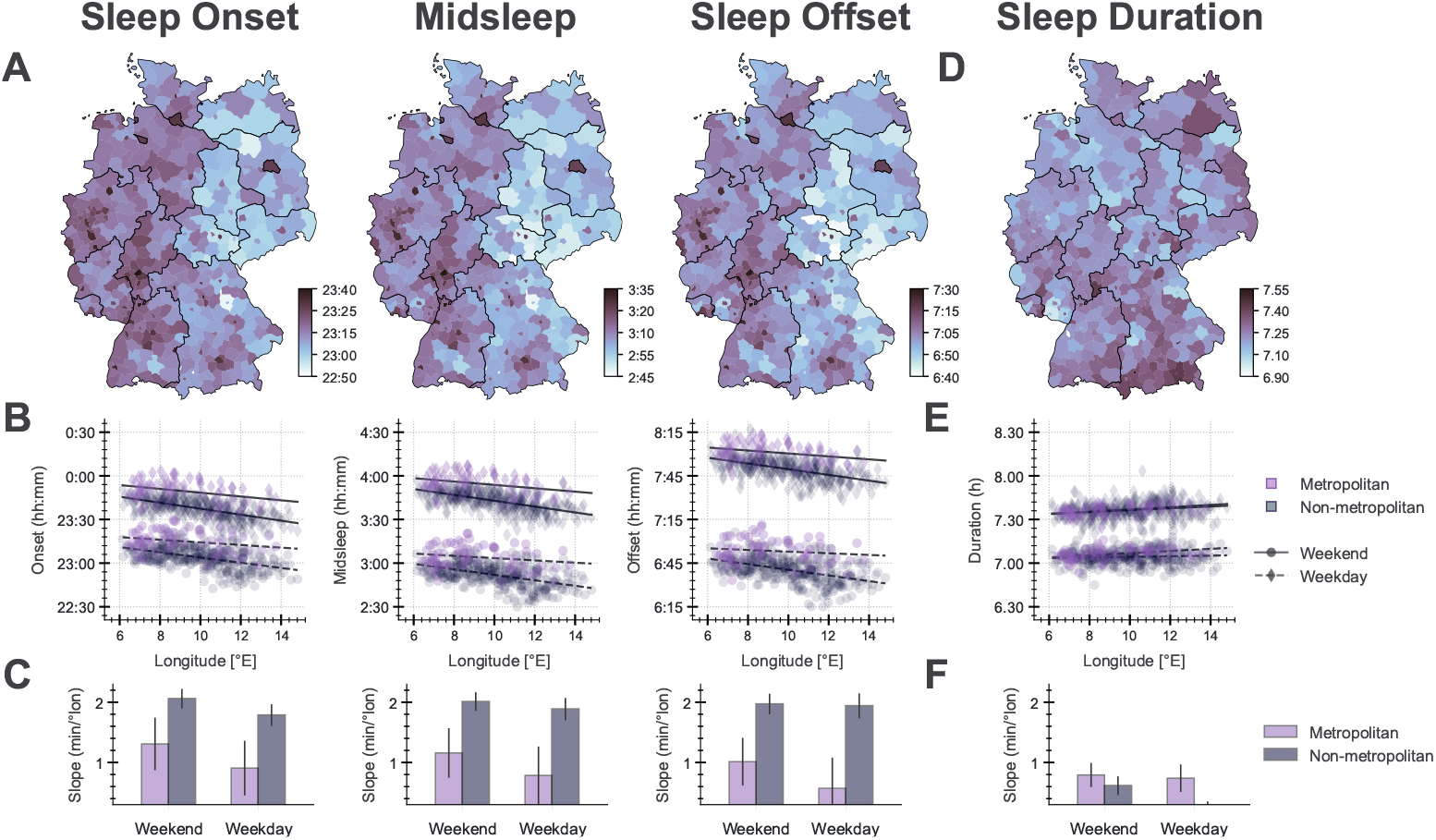
Geographic variation and longitudinal gradients in sleep timing and duration across Germany. (A) Average sleep onset, midsleep, and offset times by NUTS-3 district. (B) Associations between district longitude and sleep timing, stratified by urbanization (metropolitan: purple; non-metropolitan: gray) and day type (weekend: solid lines with circles; weekday: dashed lines with diamonds). (C) Estimated slopes (95% CI) from unadjusted linear regressions, expressed in minutes per degree longitude. (D–F) Corresponding panels for sleep duration. *Note: All regressions shown in this figure use unadjusted district-level data for visualization. Final estimates are derived from fully adjusted mixed-effects models (see Supplementary Information)*

This spatial pattern was confirmed by linear mixed-effects models controlling for demographic, geographic, and seasonal covariates (Tab. 2; Figs. S5, 6). Negative coefficients for longitude across sleep onset, midsleep, and offset indicated systematic westward delays in sleep timing with each degree of increasing westward longitude. The strongest effect emerged for weekend midsleep, with delays of 2.16 minutes per degree in non-metropolitan areas and 1.26 minutes per degree in cities. Weekday patterns were similar but weaker, with delays of 1.80 and 0.30 minutes in non-metropolitan and metropolitan regions, respectively (Tab. S3; Figs. S5, 7). Comparable gradients were observed for onset and offset. On weekends, sleep onset was delayed by 2.46 minutes and offset by 1.80 minutes per degree in non-metropolitan districts; in cities, delays were 1.50 and 0.90 minutes, respectively (Tab.2). On weekdays, the corresponding delays were 1.86 and 1.74 minutes in non-urban areas, and 0.48 and 0.12 minutes in metropolitan regions (Tab. S3).

The strength of these timing gradients varied across seasons. In non-metropolitan regions, weekend midsleep delays ranged from 2.52 minutes per degree in summer to 1.08 minutes in winter, with intermediate values in spring (1.62) and fall (2.16) (Fig. S5B–C). Seasonal variation was less pronounced in metropolitan areas and on weekdays. For example, weekday midsleep delays in cities ranged from 0.60 minutes per degree in summer to near-zero in winter. These findings suggest that both season and social constraints modulate the longitudinal gradient in sleep timing.

Across Germany’s 9-degree longitudinal span, these per-degree effects translate into substantial cumulative differences. For weekend midsleep, the east-west delay reached about 20 minutes in non-metropolitan areas and 11 minutes in cities. Weekday midsleep delays were dampened to 17 minutes and 3 minutes, respectively. Onset and offset followed similar patterns, indicating that the entire sleep episode shifts later from east to west.

Sleep duration varied less systematically with longitude than timing measures. Unlike timing, which was delayed from east to west, duration increased slightly from west to east (Fig. 1D–F). On weekends, duration increased by approximately 1.0 minute per degree of longitude in non-metropolitan areas and 0.8 minutes in metropolitan districts (Tab.2). The effect was further attenuated on weekdays, with an increase of 0.4 minutes per degree in rural regions and a nonsignificant 0.5-minute trend in cities (Tab. S3). These trends correspond to cumulative weekend differences of 9 minutes (non-metropolitan) and 7 minutes (metropolitan), and only 4–5 minutes on weekdays. Although consistent, these differences were in opposite direction, visually subtle, and an order of magnitude smaller than the timing gradients.

Overall, sleep timing delays systematically increase from east to west across Germany. These gradients are strongest on weekends and in non-metropolitan areas, and are modulated by both season and day type. By contrast, sleep duration varies only modestly across longitude, with smaller and directionally opposite effects.

### 2.2 North-South patterns in sleep — Opposing weekday and weekend trends

In addition to the prominent east-west gradient in sleep timing, we investigated variation in sleep patterns along the north-south axis. Unlike longitude, which corresponds to a relatively stable solar delay throughout the year, north–south variation reflects seasonal and latitudinal changes in solar exposure, particularly in sunrise and sunset timing. Given this complexity, we anticipated more nuanced sleep patterns associated with latitude.

While no obvious latitudinal trends were observed in initial visualizations of regional sleep maps (Fig. 1A), stratifying by day type revealed a consistent divergence between weekdays and weekends (Fig. 2A). On weekdays, participants residing farther north slept earlier than those in the south, with midsleep advancing by 0.90 minutes per degree latitude, corresponding to a 6.48-minute earlier midsleep from south to north across Germany’s 7.2° span (Tab. S3). This shift was primarily driven by earlier sleep onset (−1.14 min/°), with a smaller contribution from earlier offset (–0.66 min/°). On weekends, this pattern reversed slightly: midsleep was delayed in the north by 0.24 minutes per degree, or about 1.7 minutes in total (Tab.2). The weekend effect was primarily due to later wake times in the north (offset delay of 0.36 min/°), with a smaller effect on sleep onset (0.18 min/°). These results suggest that weekday timing shifts in northern regions reflect earlier bedtimes, while weekend shifts reflect extended morning sleep.

**Fig. 2.**
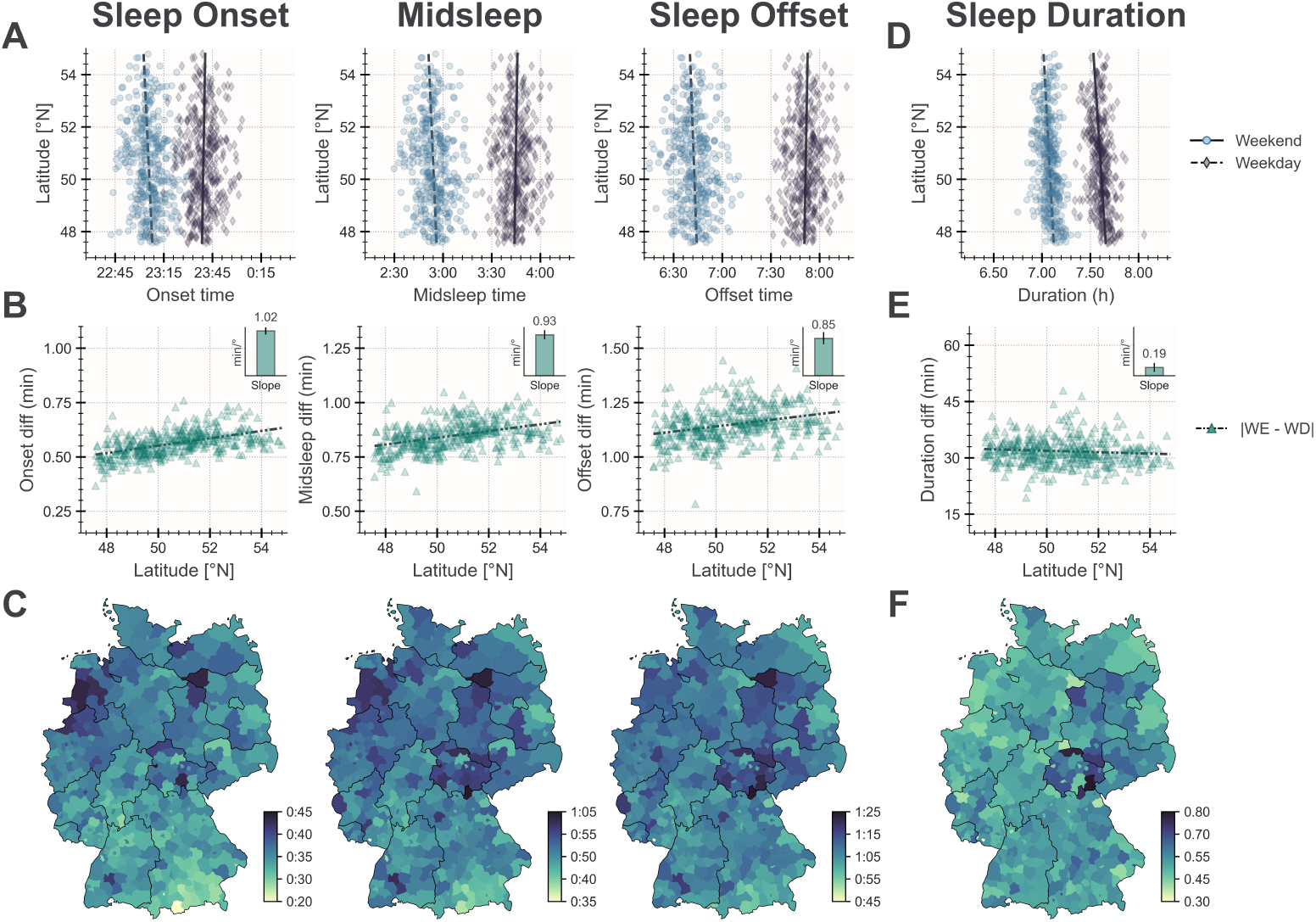
Latitudinal patterns in sleep timing, duration, and social jetlag. (A) District-level averages of sleep onset, midsleep, offset, and duration plotted against latitude, stratified by day type (weekend: blue; weekday: gray). Higher latitudes represent more northern regions. (B) Latitude-dependent differences in weekday–weekend sleep timing. Inset panels show linear regression slopes in minutes per degree latitude. (C) Maps of district-level social jetlag (weekday–weekend timing differences) indicate stronger circadian misalignment in northern and northwestern Germany. (D–F) Equivalent panels for sleep duration. *Note: All trends shown are based on unadjusted simple linear regressions using NUTS-3 district-level aggregated data. These results are intended for descriptive visualization and do not adjust for covariates or repeated measures. Fully adjusted effect estimates are reported in the main text and the Supplementary Information based on mixed-effects models*.

District-level linear regressions confirmed this divergence across all timing measures, with larger weekday–weekend differences in northern regions (Fig. 2). These differences are consistent with greater social jetlag in the north and match patterns from mixed-effects models (Tab. 2,Tab. S3). Urbanization had no significant effect on these latitudinal trends and was therefore excluded from the analysis.

Sleep duration followed a more consistent latitudinal gradient, with shorter sleep in the north across both weekdays and weekends. On weekends, individuals in northern regions slept approximately 4.1 minutes less per day than those in the south (–0.57 min/°), and 1.9 minutes less per day on weekdays (–0.27 min/°), amounting to nearly 18 minutes of cumulative weekly sleep loss (Fig.2, Tab.2, Tab. S3). This loss exceeded expectations based on sleep timing shifts alone. In fact, based on changes in sleep onset and offset times, sleep duration should have increased from south to north by approximately 19.9 minutes per week. This is because on weekdays, sleep onset advanced more than offset (1.14 vs. 0.66 min/°), and on weekends, offsets were delayed more than onsets (0.36 vs. 0.18 min/°), both of which should have yielded modest gains in sleep duration. However, models showed a total weekly sleep deficit of 17.5 minutes, rather than a gain. This discrepancy may reflect environmental, behavioral, or health-related factors at higher latitudes that suppress sleep opportunity or quality.

In summary, the north–south patterns in our dataset reflect more subtle and complex dynamics than those observed along the east–west axis. Sleep timing showed opposite trends across weekdays and weekends, while sleep duration was consistently shorter in the north. These findings suggest greater social jetlag and potential circadian misalignment among individuals living in northern Germany.

### 2.3 Seasonal trends in sleep — Sleep oscillations coincide with annual solar changes

To examine the influence of seasonal variation in light exposure on sleep behavior, we analyzed temporal patterns in sleep timing and duration across nearly three years of data. Both observed values and model-based estimates revealed consistent seasonal trends across all sleep variables, partially aligned with annual changes in solar timing (Fig. 3; Fig. S8). These oscillations were more pronounced on weekends, when external constraints such as work and school schedules were reduced. On weekdays, these patterns persisted but were consistently attenuated, reflecting the impact of social constraints on sleep behavior.

**Fig. 3.**
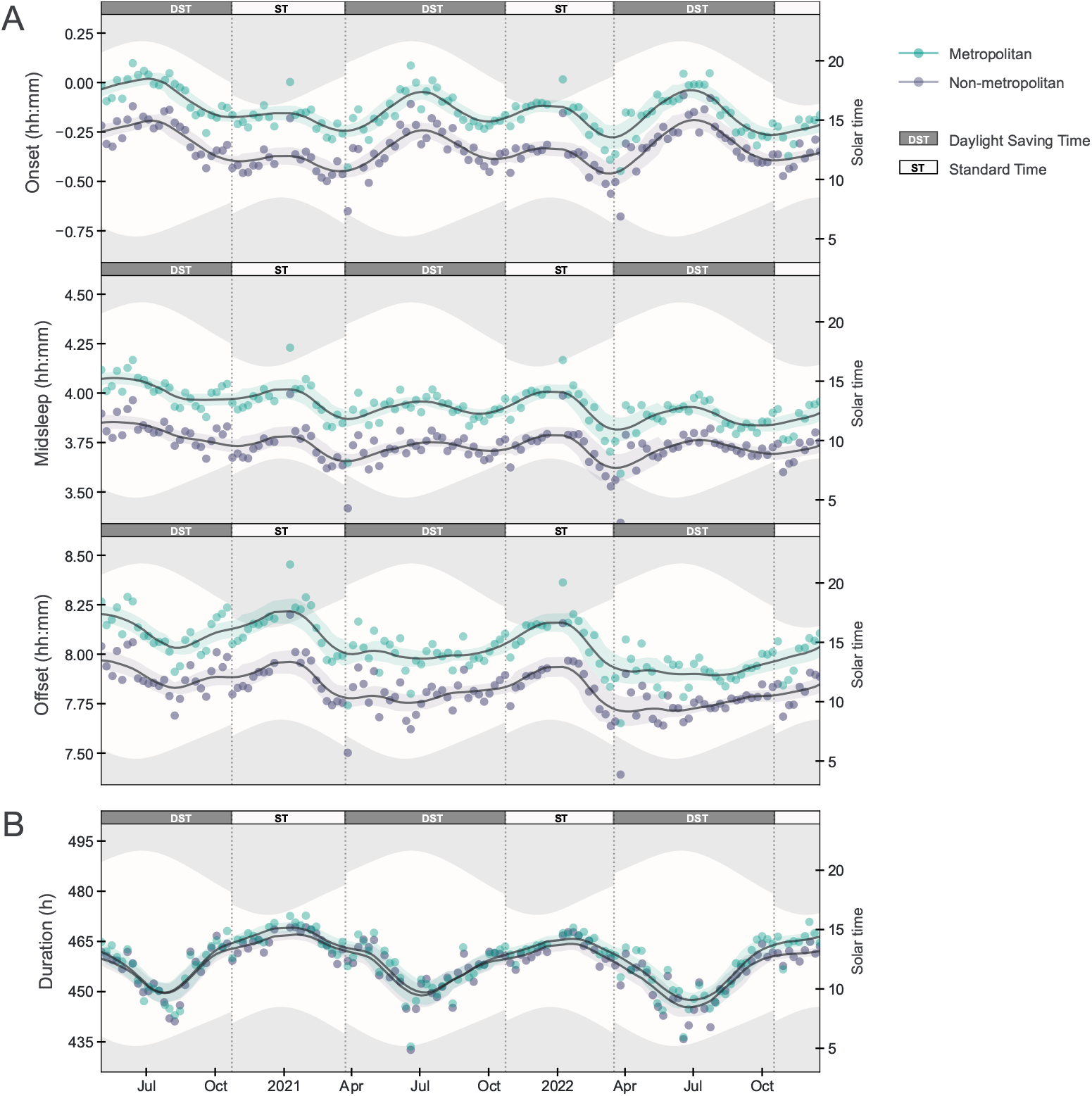
Seasonal oscillations in weekend sleep timing and duration. (A) Temporal trends in sleep onset, midsleep, offset, and duration over a 20-month period. Solid lines indicate smoothed rolling averages of aggregated weekend sleep data, stratified by urbanization (metropolitan: teal; non-metropolitan: purple). Vertical dashed lines mark daylight saving time (DST) transitions; shaded bands indicate average photoperiod, bounded by mean daily sunrise and sunset times. Sundays falling on DST transitions and weekends during the winter holiday period (December 24–January 3) were excluded. (B) Annual variation in sleep duration. Duration peaks in winter (standard time) and reaches a minimum in summer (DST), consistent with seasonal variation in both light exposure and daily routines. This seasonal pattern is more pronounced on weekends than weekdays (see Fig. S8), indicating that social constraints dampen sleep oscillations during the workweek. *Note: DST and standard time (ST) labels are used in place of seasonal labels to reduce visual clutter and improve interpretability*.

Sleep duration showed the strongest seasonal modulation, forming a sinusoidal waveform that peaked in winter and dipped in summer (Fig. 3B). Mixed-effects model estimates confirmed this pattern: individuals slept 24.7 minutes longer in winter compared with summer on weekends, and 11.6 minutes longer on weekdays (Tab. 2; Tab. S3). These differences amount to nearly two hours of cumulative sleep loss across a summer week compared to winter, consistent with previous findings [24–29]. The reduced weekday amplitude was due primarily to a shallower summer trough, while the winter peak was relatively preserved (Fig. S8B).

Sleep offset followed a similarly consistent sinusoidal rhythm, with later wake times in winter and earlier wake times in summer. This pattern closely mirrored seasonal shifts in sunrise (Fig. 3A, bottom row). Model estimates indicated that wake time was delayed by 21.8 minutes in winter relative to summer on weekends, and by 15.9 minutes on weekdays (Tab. 2; Tab. S3). As with sleep duration, weekday effects were smaller but followed the same direction (Fig. S8).

In contrast, sleep onset exhibited a biannual pattern, with delays in both summer and winter and earlier bedtimes in spring and fall (Fig. 3A, top row). Model estimates showed that, on weekends, sleep onset was delayed by 1.44 minutes in winter and 2.64 minutes in summer relative to fall (Tab. 2). On weekdays, the same seasonal effects were reduced or nonsignificant (winter: –3.42 min; summer: +0.24 min), indicating that this rhythm was notably dampened by weekday constraints (Fig. S8; Tab. S3). The distinct shape of this waveform differentiates sleep onset from other outcomes, which more closely follow a single annual cycle.

Midsleep displayed an asymmetric biannual rhythm shaped jointly by onset and offset timing (Fig. 3A, middle row). On weekends, midsleep was delayed by 21.5 minutes in winter compared to summer; on weekdays, the delay was 11.3 minutes (Tab. 2; Tab. S3). The shape of the midsleep waveform reflected the combined influence of both onset and offset times. As with other outcomes, weekday constraints reduced the seasonal amplitude, mainly by flattening the summer advance rather than shifting the winter peak (Fig. S8).

Together, these results show that sleep timing and duration systematically vary with season, with more pronounced oscillations on weekends than weekdays. The amplitude and shape of these rhythms differ by sleep outcome, with sleep duration and offset following sinusoidal patterns, sleep onset exhibiting a biannual curve, and midsleep reflecting a composite profile.

## 3 Discussion

This study reveals robust spatial and temporal patterns in human sleep behavior, shaped by the interaction between solar timing and social constraints. The clearest and most consistent finding was a longitudinal gradient in sleep timing: across all timing outcomes, individuals in western Germany slept later than those in the east, despite residing in the same time zone. This east–west delay reflects a westward shift in solar time relative to local time, persisting even after adjusting for demographic, seasonal, and geographic covariates. These results align with previous studies based on self-reports and chronotype assessments [17–20, 30, 31] and confirm that natural solar cues continue to influence population-level sleep timing, even in highly industrialized settings.

The strength of this gradient was moderated by both urbanization and day type, reflecting the role of artificial lighting and structured schedules in shaping sleep timing. In metropolitan regions, the east–west difference was reduced by nearly half, suggesting that city dwellers are less entrained to solar time. On weekdays, the gradient was similarly attenuated, consistent with stronger alignment to work and school schedules. These patterns indicate that modern social structures modulate, but do not fully override, circadian alignment with solar cues. Seasonal variation in the gradient further supports this interpretation: the slope was steeper in summer and fall, when daylight is longer and behavioral flexibility may be greater, and flatter in winter and spring.

By contrast, sleep duration varied only modestly with longitude. Although statistically significant due to the large sample size, the effect was roughly one-tenth the size of the timing gradient and even smaller on weekdays. This suggests that while solar timing exerts a strong influence on when people sleep, it has far less impact on how long they sleep.

Compared to the robust east–west gradient in sleep timing, north–south patterns were subtler and more complex. They were characterized by bidirectional effects consistent with social jetlag, likely reflecting both solar timing and regional social norms. On weekdays, earlier bedtimes and wake times in northern Germany may stem from earlier work or school start times, or other cultural factors. This shift was primarily driven by earlier sleep onset, suggesting that social constraints played a key role. On weekends, this pattern reversed: people in southern Germany slept earlier than those in the north, primarily due to later wake times in northern regions. This may reflect weaker morning light exposure in the north and greater circadian entrainment in the south, where solar intensity is higher. Together, these opposing weekday and weekend trends point to greater misalignment between biological and social timing in northern Germany.

Unlike sleep timing, sleep duration declined consistently from south to north, across both weekdays and weekends. This amounted to nearly 18 minutes of weekly sleep loss in the north, which contradicts expectations based on sleep timing alone. Model-estimated changes in onset and offset predicted a weekly gain of about 20 minutes, due to earlier bedtimes on weekdays and later wake times on weekends. The observed loss, despite earlier bedtimes on weekdays and later wake times on weekends, suggests lower sleep efficiency or reduced sleep opportunity. Potential contributors include reduced daylight exposure, increased artificial light in the evening, or other socio-environmental stressors.

These findings indicate that northern populations may face a dual burden: greater misalignment between social and biological timing, and shorter, potentially lower-quality sleep. While modest in magnitude, these effects were consistent, underscoring the need to consider both solar cues and regional social context in sleep research.

Seasonal variation added a temporal layer to these spatial effects. Sleep timing and duration followed systematic annual cycles, with later and longer sleep in winter and shorter, earlier sleep in summer. These oscillations were more pronounced on weekends, suggesting that weekday schedules suppress seasonal adaptation. Among all outcomes, sleep offset tracked solar timing most closely, while onset followed a biannual rhythm, with delays in both summer and winter and earlier bedtimes in spring and fall. This may reflect competing effects: extended daylight delays sleep in summer, while increased evening activity and artificial light contribute to winter delays. Midsleep, shaped by both onset and offset, captured this composite seasonal response.

Together, these findings reinforce the dual role of solar and social time in shaping sleep behavior. Solar cues anchor underlying circadian rhythms, especially in rural regions and on unconstrained days. Social obligations modulate how these rhythms are expressed in daily life. By integrating large-scale wearable data with high-resolution spatiotemporal modeling, this study quantifies how geography, season, and social context jointly influence population-level sleep.

### 3.1 Health Implications and Policy Considerations

The persistence of east–west differences in sleep timing, even under artificial lighting and modern social constraints, underscores the continuing influence of solar cues on human circadian rhythms. The seasonal amplification of these gradients suggests that solar entrainment is strongest during spring and summer, when daylight exposure is greatest. However, the resulting misalignment between solar and social time, particularly in western and northern regions, may contribute to circadian disruption and increased health risk.

Regions with delayed sleep timing, such as western Germany, or shorter sleep duration, as observed in northern Germany, may experience elevated health disparities due to chronic circadian misalignment. Social jetlag, which was most pronounced in northern areas, has been consistently linked to adverse health outcomes, including metabolic dysfunction, cardiovascular disease, and reduced overall well-being [32, 33]. This misalignment has also been linked to higher rates of workplace accidents and motor vehicle collisions, particularly in areas where solar and social timing are most misaligned [34, 35].

These findings point to the need for public health strategies that account for geographic and seasonal variation in sleep behavior. Policies should aim to mitigate the disruptive effects of daylight saving time (DST), which increases solar-social misalignment, and promote schedules that are better synchronized with human circadian physiology. While modern life has reduced natural entrainment, solar time remains a key reference point for biological rhythms. Adapting social structures to better reflect this alignment could improve sleep health and reduce regionally patterned disparities.

## 4 Limitations

Despite the strengths of this study, particularly its large-scale dataset, high temporal resolution, and objective sleep measures, several limitations must be acknowledged. These relate primarily to sampling bias, methodological heterogeneity, and the influence of the COVID-19 pandemic, all of which may affect the generalizability of the findings.

First, data collection occurred during the COVID-19 pandemic, a period marked by widespread disruptions to daily life and sleep behavior. Approximately 40% of the German population reported significant changes in their sleep patterns, including longer and later sleep schedules, reduced social jetlag, and declines in sleep quality [27, 36–40]. Although relaxed social constraints during the pandemic may have amplified solar influences on weekday sleep timing, the spatial patterns observed here are broadly consistent with pre-pandemic studies [17]. Still, the extent to which these results generalize to post-pandemic conditions remains uncertain.

Second, the sample is not fully representative of the German population. Participation skewed toward middle-aged adults, with limited representation of adolescents (under 20 years) and a decline in participation among those over 60 (Fig. S2A). This likely reflects both age-related differences in digital technology adoption and the study’s minimum age requirement (16+ years) [41]. In addition, the sample was 52% male and likely biased towards higher socioeconomic status and health-conscious individuals, as fitness tracker ownership is more common among affluent and physically active populations [42]. Although mixed-model analyses indicated minimal impact of demographic factors on sleep timing and duration (Fig. S1), future research should prioritize more inclusive sampling, particularly among adolescents, elderly individuals, and socioeconomically diverse groups.

Third, the study relied on data from multiple consumer-grade wearable devices, with no control over hardware specifications, firmware versions, or proprietary sleep detection algorithms. These algorithms typically use accelerometry and heart rate variability, but differences across manufacturers may affect sleep classification. Notably, data from Apple devices (representing about one-third of participants) were excluded due to a software update in October 2021 that rendered sleep measures incompatible with earlier data. While many wearables show reasonable agreement with polysomnography or actigraphy for sleep–wake detection [43], device-specific differences limit cross-device comparability. To address this, the present study focused on consistently estimated variables such as sleep onset, offset, and midsleep, and excluded measures related to sleep architecture or staging. Even so, device variability remains a concern and highlights the need for standardized validation protocols and harmonized processing pipelines in wearable-based sleep research.

Despite these limitations, the findings provide robust evidence of systematic spatiotemporal variation in sleep behavior and reinforce the dual influence of solar cues and social timing. Future studies should aim to replicate these results in more diverse populations, develop validated methods for integrating wearable data, and further explore environmental and social determinants of sleep behavior.

## 5 Conclusion

This study demonstrates the potential of passive sensing technologies to advance public health monitoring by enabling the large-scale, continuous, and objective measurement of sleep behavior with high temporal resolution. Compared to traditional surveys and self-reports, wearable-based data provide non-invasive, real-time insights that capture longitudinal patterns and environmental influences with greater precision. To our knowledge, this dataset represents the largest study of its kind to objectively examine how sleep timing and duration vary across geography and season. By leveraging high-resolution wearable data, this research contributes to the growing field of digital epidemiology and supports the use of passive sensing in population-level sleep and circadian studies. Our findings reinforce the ongoing influence of solar time on human sleep, even in the context of artificial lighting and structured daily schedules.

The consistent misalignment between biological and social time, particularly in western and northern Germany, highlights the need for regionally informed public health strategies. Policies that promote better alignment between societal timing and natural circadian rhythms—such as delaying school start times, offering flexible work schedules, or eliminating daylight saving time—may help mitigate circadian disruption and improve sleep health. Ultimately, this work underscores the importance of integrating both solar and social timing factors into sleep science and public health research. These insights can support targeted interventions designed to foster healthier, more synchronized sleep patterns across diverse populations.

## 6 Methods

### 6.1 Data collection

Sleep data were collected via passive sensors in wearable devices as part of the Robert Koch Institute’s Corona Data Donation project (*Corona-Datenspende*), launched in April 2020. By downloading the *Corona-Datenspende* app, consenting to the data protection declaration, and linking a supported consumer smartwatch or fitness tracking device, German residents aged 16 years and older were able to provide daily physiological and behavioral measurements, including heart rate, activity, and sleep metrics. Supported device platforms included Fitbit, Garmin, Google Fit, Huawei Health, Oura, Polar, Amazfit, Withings and Apple Health. Each day, device data were acquired, de-identified, and harmonized into a composite data set by a wearable data connectivity platform developed by Thryve (Germany, https://thryve.health). To participate, users were required to provide either their full postal code (five digits) or a partial code (first three digits), depending on the date of enrollment, enabling regional analysis of sleep patterns. They also could optionally report gender, age, height, and weight (in 5-unit intervals) for cohort analyses. Body mass index (BMI) was estimated when both height and weight were provided, by centering the reported values within their respective 5-unit bins. For the present analysis, sleep data provided between May 1, 2020 and December 31, 2022 were considered.

### 6.2 Ethical considerations

Participation in the Corona Data Donation Project was voluntary and self-recruited, open to all German residents aged 16 and older. Participants provided informed consent electronically via the *Corona-Datenspende* smartphone app. All participant data were de-identified and stored pseudonymously using a randomly generated unique identifier. No identifying information (e.g., names, addresses, birth dates) was collected or stored at any time. The study was conducted in full compliance with the European Union’s General Data Protection Regulation (GDPR), ensuring strict standards for personal data protection. The project was reviewed and approved by the Data Privacy Officer at the Robert Koch Institute (internal operation number 2021-009), in agreement with the Federal Commissioner for Data Protection and Freedom of Information (BfDI), Germany’s highest independent authority for data protection and freedom of information. Ethical approval for the Corona Data Donation project was also obtained from the ethics board at the University of Erfurt (approval number 20220414).

### 6.3 Statistical analysis

Statistical analyses were conducted using linear mixed-effects models to estimate the influence of geographic, temporal, demographic, and environmental variables on sleep timing and duration. Models included participant-level random intercepts to account for repeated measures and were adjusted for relevant covariates. Full modeling specifications, preprocessing steps, and variable definitions are provided in the Supplementary Information.

## 8 Acknowledgments

We are deeply grateful to the participants of the Corona Data Donation Project, whose contributions made this research possible. We thank Paul Burggraf for his technical support during the data collection process, and Claudia Enge and Lorenz Wascher 15 for their ongoing guidance on data privacy and protection matters. We also acknowledge Jakob Kolb and David Hinrichs for their valuable assistance with data curation, infrastructure, and administrative coordination. Finally, we thank Marc Wiedermann for his foundational contributions to the development of the data donation platform and the early stages of this project.

## 9 Data availability

The data used in this study are available upon reasonable request. Due to privacy and data protection regulations, access requires registration and approval by the Data Privacy Officer at the Robert Koch Institute. Requests should be directed to Prof. Dirk Brockmann

## 10 Code availability

All custom code used for data preprocessing, analysis, and figure generation is available upon request. Code can be shared under a data use agreement to ensure alignment with data privacy policies and ethical guidelines. Interested researchers should contact Prof. Dirk Brockmann for access.

## 11 Author contributions

AR led the data analysis, validation, visualization, and software development, and was responsible for drafting and revising the manuscript. ECW and DB contributed to the conceptualization and supervision of the project, with ECW focusing on sleep-related analyses and DB overseeing broader study coordination. Methodology development was carried out by AR, HS, and ECW. Data curation and investigation were performed by AR and HS. Funding acquisition and project administration were managed by DB and BFM, who also played key roles in the design and implementation of the Corona Data Donation platform. Resources were provided by DB. All authors contributed to reviewing and editing the manuscript.

