## Supplementary Information for "Large-scale wearable data reveal spatiotemporal organization of annual sleep patterns"

.

†Present address(es): Benjamin F. Maier, Freelance Researcher, Berlin, Germany.

Contributing authors:;

### Supplementary Methods

This Supplementary Data file provides additional methodological detail and extended analyses supporting the main text.

#### Sleep Variables

Sleep patterns were assessed on the basis of four core variables: sleep duration, sleep onset, sleep offset, and midsleep time. Midsleep, calculated as the midpoint between onset and offset, served as the primary sleep timing outcome. It provides a robust and concise measure of sleep timing, that, unlike onset or offset, is independent of duration yet remains sensitive to changes in both [1]. Midsleep is also strongly correlated with dim light melatonin onset (DLMO), the gold-standard circadian biomarker, making it a validated proxy for internal circadian phase and chronotype in large-scale field studies [2, 3].

Wearable-derived metrics characterize the main sleep episode each day, defined as the longest continuous period of sleep detected within a 24-hour cycle. Sleep onset and offset refer to the start and end of this episode, while duration reflects the elapsed time between them, excluding interim wake periods.

To account for variation in sleep behavior due to social constraints (e.g., work or school), sleep variables were analyzed separately for weekdays and weekends. Weekend nights were prioritized in primary analyses, as they more closely reflect individuals' endogenous sleep patterns in the relative absence of external obligations [4].

Timestamps were converted to Central European Time (CET) using transmitted time zone offset values. Sleep episodes were assigned to the calendar day of sleep offset. For example, a "Saturday sleep value" corresponds to the primary sleep episode occurring from Friday night to Saturday morning. Sleep timing variables were recoded as decimal hours relative to midnight (e.g., -1.5 = 10:30 PM), and sleep duration was expressed in minutes.

For information on device-specific variability and limitations in sleep detection, see the Limitations.

##### 0.1 Inclusion criteria

To be included in the analysis, participants were required to have at least 20 valid weekdays and 10 valid weekend days of sleep data during the study period. Sleep episodes were assigned to the day of offset, with start times permitted to fall between noon of the previous day and noon of day of wake-up. To remove extreme or likely unreliable data, participants with an average sleep duration of less than 5 hours or more than 12 hours were excluded. These thresholds are based on sleep research guidelines that identify such durations as uncommon or indicative of measurement error [5]. In addition, individuals were excluded if any daily sleep values exceeded 24 hours, or if sleep offset occurred before midnight on the day of wake or after midnight the following day. Such values are physiologically implausible and typically result from device synchronization errors. Permissible ranges were thus:

- Duration: 0 to 24 hours

- Onset: 12:00 (previous day) to 12:00 (wake-up day)
- Offset: 00:00 to 24:00 (wake-up day)

Participants contributing data from more than one device source were excluded to avoid inconsistencies. We also removed data recorded outside the Central European Time zone, which could reflect travel-related disturbances or incorrect device time settings. Due to a significant change in Apple's proprietary algorithms used for sleep quantification introduced in October 2021, all Apple Health data were excluded from the analysis to maintain consistency across individuals and devices.

To minimize the influence of atypical events on population-level sleep trends, the following days were excluded from all analyses:

- National and regional public holidays falling on weekdays
- December 24 to January 3
- Sundays of DST transitions and the five days following each transition [6, 7]
- Outlier days with population-level sleep metrics exceeding 1.5 times the interquartile range, calculated separately for weekends and weekdays

#### 0.2 Spatiotemporal variables

Self-reported postal codes were geocoded to latitude and longitude using the GeoNames geographical database (https://download.geonames.org/export/zip, accessed January 2022). Postal codes were also mapped to districts, known as "Kreise" or "kreisfreie Städte", according to the Level 3 NUTS3 (Nomenclature of Territorial Units for Statistics) classification, using correspondence tables provided by Eurostat (https://ec.europa.eu/eurostat/web/nuts/correspondence-tables/postcodes-and-nuts, accessed January 2022).

These geographic coordinates were then used to derive local daily sunset and sunrise times based on the sunrise equation using the Python package *suntime* [8].

Solar seasons were used, as opposed to meteorological or astronomical seasons, to provides balanced season lengths centered around the equinoxes and solstices, which helps align seasonal analyses with meaningful changes in light exposure. Seasons were defined as follows:

- Spring: February 5 – May 6
- Summer: May 7 – August 6
- Fall: August 7 – November 5
- Winter: November 6 – February 4

This seasonal classification is particularly relevant for sleep research, as it captures changes in daylight duration that influence circadian timing and behavior.

Urbanization classifications were based on the "siedlungsstrukturelle Kreistypen" (settlement-structural district types) provided by the INKAR database from the German Federal Institute for Research on Building, Urban Affairs and Spatial Development (https://www.inkar.de, accessed October 2021). Each of the 401 NUTS-3 regions was assigned to one of four structural types based on population size and settlement density:

1. Independent cities ("kreisfreie Städte") with at least 100,000 inhabitants
2. Urban districts
3. Rural districts with densification approaches
4. Sparsely populated rural districts

For analysis purposes, we classified all independent urban districts (type 1) as "metropolitan," and grouped types 2–4 into a single "non-metropolitan" category.

#### 0.3 Data analysis

Data were processed in Python (version 3.12.8) using the following packages: `pandas` (version 2.2.3) and `numpy` (version 2.2.4) for data manipulation and aggregation; `matplotlib` (version 3.10.1), `seaborn` (version 0.13.2), and `geopandas` (version 1.0.1) for visualization [9**?** –12]. Local sunrise and sunset times were calculated using the `suntime` package (version 1.2.5) based on geocoded coordinates [8].

All mixed-effects models were estimated in R (version 4.1.2) using `lme4` (version 1.1.36) and `lmerTest` (version 3.1.3) for model fitting [**? ?** ]. Marginal means and contrasts were derived using `emmeans` (version 1.10.5) and visualized using `ggplot2` (version 3.5.1) and `ggforestplot` (version 0.1.0) [**? ? ?** ]. Data wrangling was performed with `dplyr` (version 1.1.4), and model diagnostics with `performance` (version 0.13.0) [**? ?** ].

All averages (except daily means) were computed as participant-level means: data were first averaged within individuals before being aggregated across subgroups (e.g., age, region, or season). Clock time was used throughout the analyses, defined in local time (CET/CEST), consistent with the time displayed on participants' devices. As a result, differences between standard time and daylight saving time reflect changes in behavior relative to the civil clock rather than to solar time.

To examine large-scale spatial trends, sleep variables were aggregated to the district level and regressed on geographic coordinates (longitude and latitude). These simple linear regressions, performed separately by day type and urbanization level, complemented the mixed-effects models by illustrating regional patterns while minimizing individual-level noise.

#### 0.4 Data modeling

We adopted a hierarchical linear mixed-effects modeling approach to assess the influence of demographic and spatiotemporal variables on sleep outcomes. Given the nested structure of the data, daily observations were grouped within individual users as random intercepts to control for interindividual variability and temporal autocorrelation. Sensitivity analyses also tested a random intercept for day, but it did not substantially alter fixed effects and was excluded from the final model for efficiency. The final model structure was selected based on a combination of prior knowledge and exploratory modeling using the primary outcome (weekend midsleep) and the Akaike Information Criterion (AIC) for model selection.

Independent variables were kept to a minimum, and interactions were limited to second-order to facilitate interpretability. The model was specified as:

$$
\begin{aligned}
y_{ij} = \beta_0 &+ \beta_1(\text{age}_{ij}) + \beta_2(\text{gender}_{ij}) + \beta_3(\text{BMI}_{ij}) + \beta_4(\text{urbanization}_{ij}) + \beta_5(\text{DST}_{ij}) \\
&+ \beta_6(\text{season}_{ij}) + \beta_7(\text{longitude}_{ij}) + \beta_8(\text{latitude}_{ij}) + \beta_9(\text{sunrise}_{ij}) \\
&+ \beta_{10}(\text{longitude}_{ij} \times \text{urbanization}_{ij}) + \beta_{11}(\text{longitude}_{ij} \times \text{season}_{ij}) \\
&+ \beta_{12}(\text{sunrise}_{ij} \times \text{season}_{ij}) \\
&+ u_{0i} + \epsilon_{ij},
\end{aligned}
$$

$$
u_{0i} \sim N(0, \sigma_u^2), \quad \epsilon_{ij} \sim N(0, \sigma^2)
$$

- $y_{ij}$ represents the sleep outcome for the $j$-th observation of the $i$-th individual.
- $u_{0i}$ is the random intercept for user $i$.
- $\epsilon_{ij}$ is the residual error.

To improve interpretability and reduce multicollinearity, continuous predictors were centered prior to model fitting. Specifically, age and BMI were centered around their sample means; longitude and latitude were centered on the geographic midpoint of Germany (10.5°E, 51°N); and sunrise time was centered on the average local sunrise time at this midpoint on the fall equinox (approximately 7:06 CET). Centering allowed regression coefficients to be interpreted as the expected change in the outcome variable for a one-unit increase in the predictor, relative to a geographically and demographically representative reference individual. Model fit was evaluated using marginal and conditional $R^2$ values, calculated via the Nakagawa method [**?** ], which capture variance explained by fixed effects alone and by the full model, respectively. For midsleep timing, the model explained 47% of the total variance (conditional $R^2$), with fixed effects accounting for 2%.

Coefficients from mixed-effects models were estimated in decimal hours and converted to minutes for presentation in the main text and Supplement.

Weather and device brand variables were excluded due to limited explanatory value or instability in estimates [**?** ].

Unless stated otherwise, model predictions represent a reference individual: female, non-metropolitan, mean age (48.6 years), mean BMI (26.3 kg/m$^2$), residing at the geographic midpoint of Germany, during fall solar season and under standard time. Because the reference sunrise time does not reflect seasonal extremes, intercepts for seasonal model predictions were manually adjusted for visualization purposes only, using average sunrise times at the geographic midpoint of Germany on the respective solstices and equinoxes: 05:04 (summer), 06:22 (spring), and 08:19 (winter). These adjustments did not affect any reported model results or statistical estimates.

#### 0.5 Sensitivity Analyses

To assess the robustness of our findings, we conducted a series of sensitivity checks examining the effects of model structure, demographic sampling, and different parameterizations of sunrise timing. These analyses confirmed that the core patterns reported remained consistent regardless of modeling choices.

Incorporating demographic covariates such as age, gender, and BMI did not materially alter the results, although it improved model performance and was retained for completeness. Similarly, including a day-level random intercept had minimal influence on estimated effects and was excluded from final models to preserve computational efficiency.

To test the role of seasonal light variation, we compared models using a fixed sunrise reference point to those using seasonally adjusted values. Both approaches yielded consistent results, supporting the robustness of the sunrise variable. Additional analyses decomposing sunrise into daily (within-person) and regional (between-person) components showed that short-term solar changes were the primary drivers of sleep timing, while longer-term regional differences played a lesser role.

Finally, we tested whether the effect of sunrise differed across regions. While the relationship between daily sunrise and sleep timing was moderated by east–west location, regional differences did not meaningfully alter the broader trends.

Together, these analyses reinforce the reliability of the findings. The observed spatial and seasonal patterns in sleep remained consistent across a range of analytical specifications, underscoring the persistent influence of both solar cues and social timing structures on human sleep behavior.

### Supplementary Figures

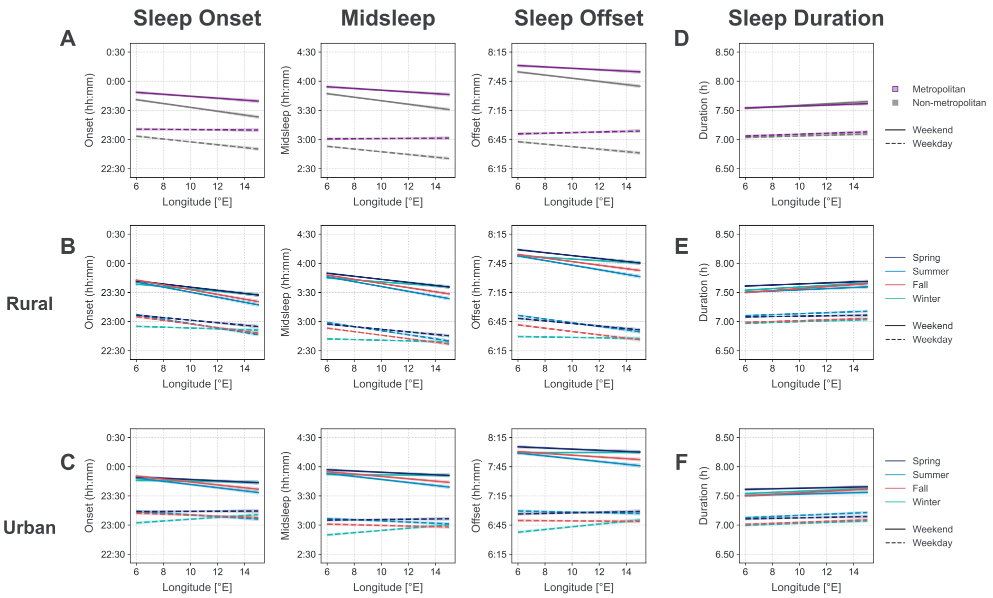


**Fig. 1**:
**Figure S1** — Distribution of sleep timing and duration by day type. Histograms of sleep onset, midsleep, offset, and duration are shown separately for weekdays and weekends. Red dashed lines indicate the sample mean for each distribution, with mean and standard deviation reported in the legend. Weekend distributions are shifted later in time and exhibit greater variability for all timing metrics. Sleep duration is also longer on weekends, reflecting the reduction of social constraints on non-workdays. Corresponding summary statistics are provided in Table 1.

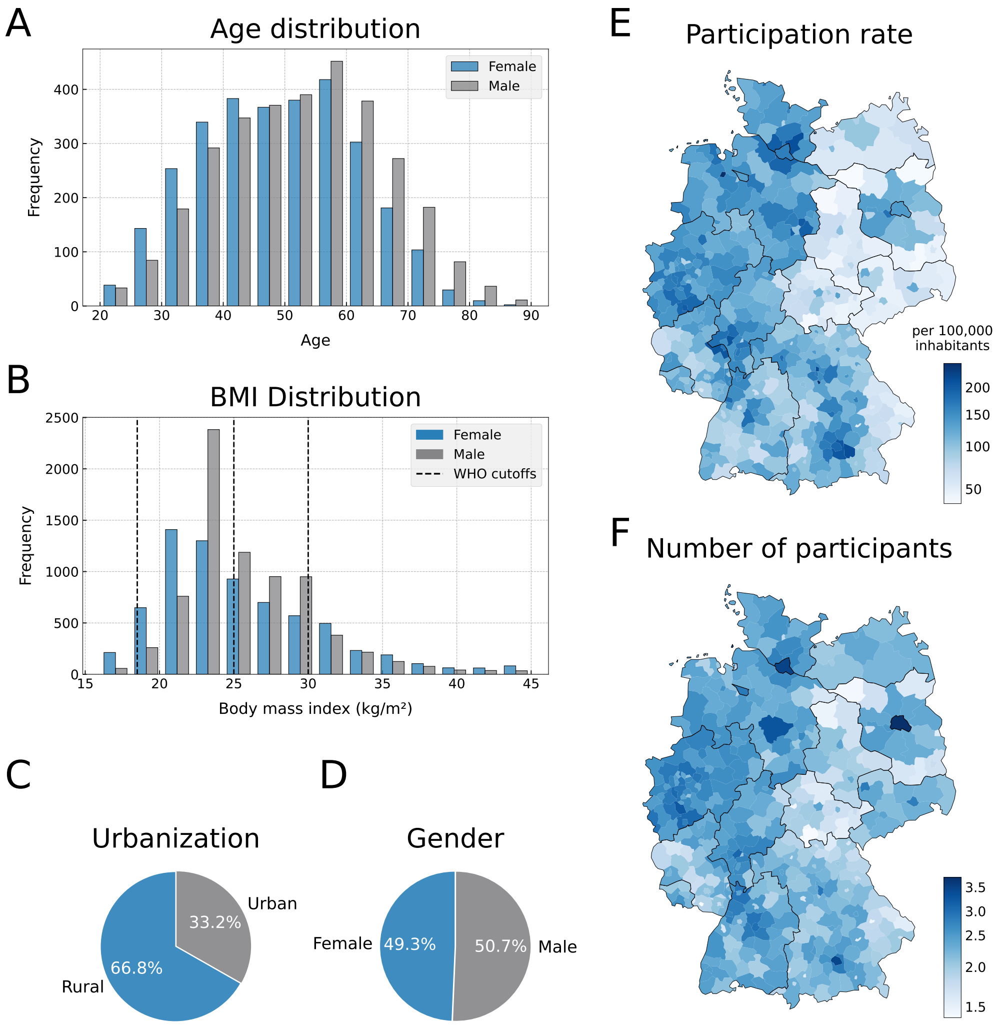


**Figure S2** — Demographic and geographic characteristics of the study population. (A) Age distribution stratified by gender. (B) Body mass index (BMI) distribution; dashed lines indicate WHO thresholds for underweight (<18.5), normal weight (18.5–24.9), overweight (25–29.9), and obese (≥30). (C) Proportion of participants classified as metropolitan vs. non-metropolitan according to district classification (see Methods for details). (D) Gender distribution of the sample. (E) Regional participation rate expressed as the number of participants per 100,000 inhabitants. (F) Absolute number of participants per district, displayed on a logarithmic scale. *Note: Panels A, B, and D are based on the subsample with self-reported demographic data (N = 59,772). Panels C, E, and F are based on ZIP code information available for the full analytic sample (N = 105,741).*

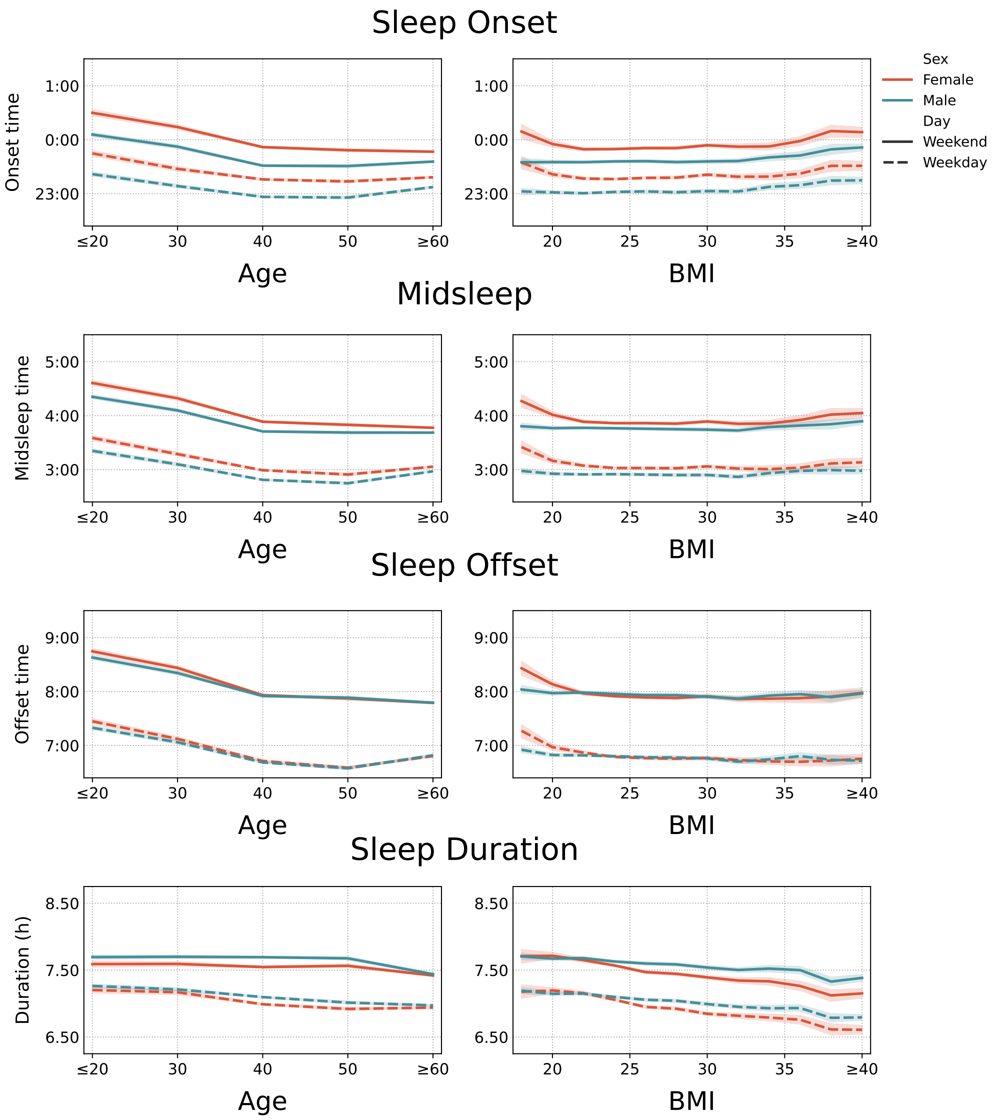


**Figure S3** — Sleep timing and duration by age and BMI. (A) Sleep onset, midsleep, offset, and duration stratified by age group. (B) Equivalent metrics stratified by body mass index (BMI). Results are shown separately by gender (female: blue, male: gray) and day type (solid lines: weekends; dashed lines: weekdays). Across both age and BMI, sleep timing shifted earlier and duration shortened with increasing age or BMI. These trends were consistent across sexes and between weekdays and weekends, with wider variability observed on weekends.

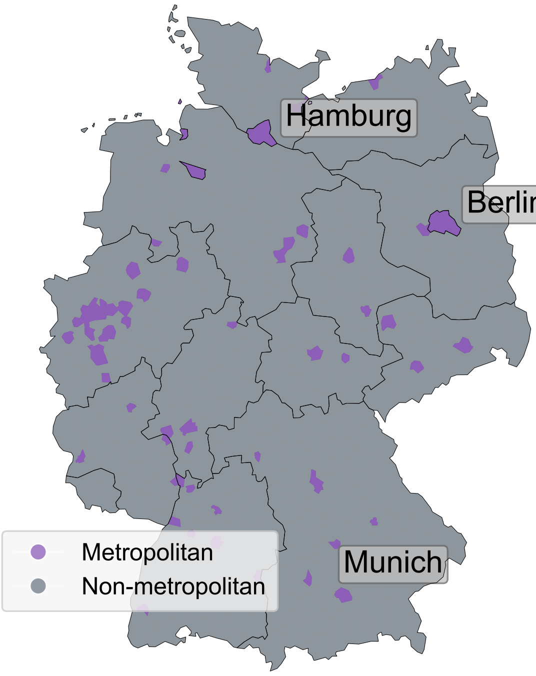


**Figure S4** — Urbanization classification of German districts. Map of NUTS-3 districts categorized by urbanization level based on classifications from the German Federal Institute for Research on Building, Urban Affairs and Spatial Development (INKAR). Metropolitan districts (purple) represent independent cities with populations exceeding 100,000. All other regions (gray), including urbanized rural districts and sparsely populated areas, are classified as non-metropolitan. Major cities (Berlin, Hamburg, Munich) are labeled for reference. This binary classification was applied in all urbanization-stratified analyses. See Methods for details on the original classification scheme and criteria.

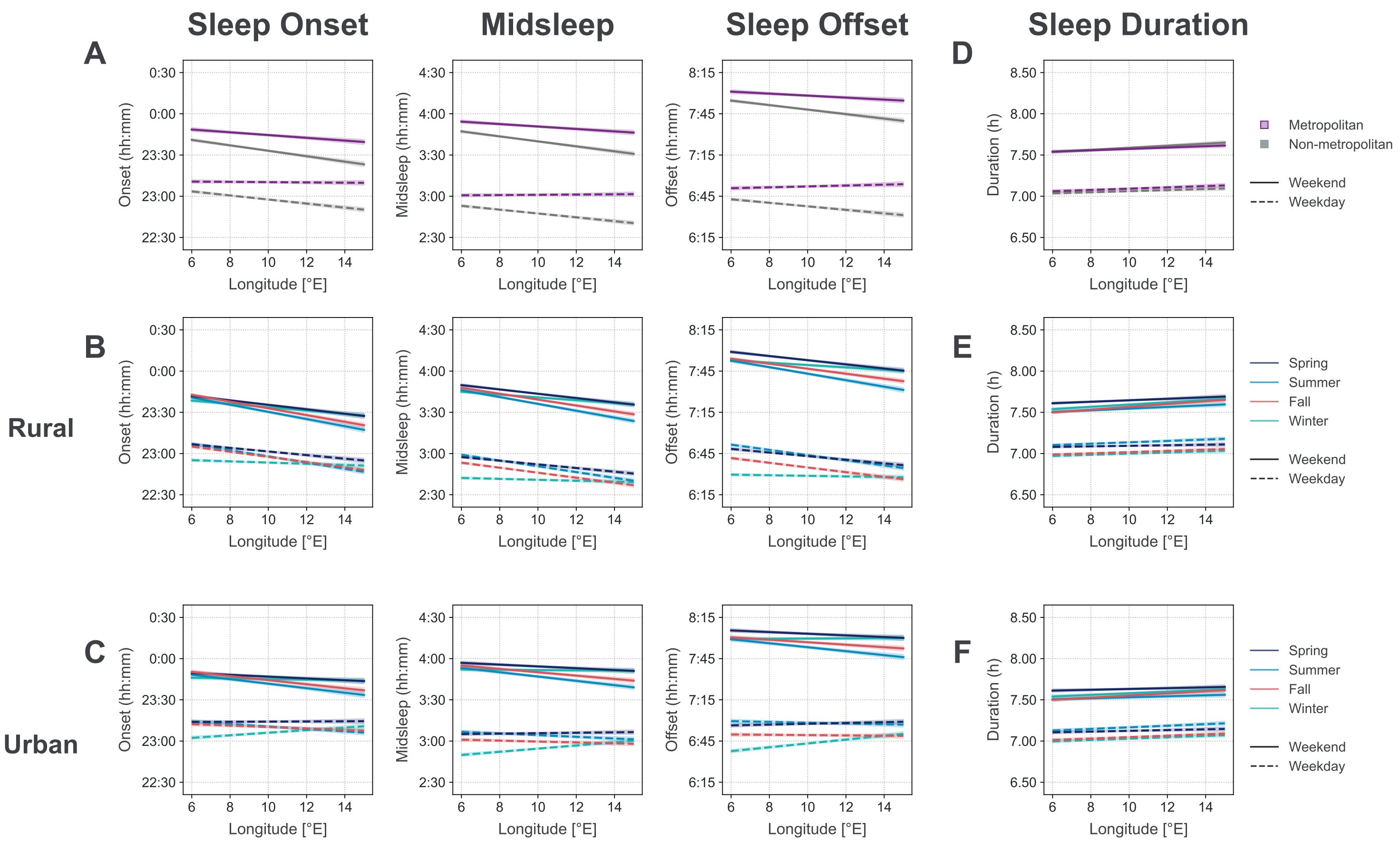


**Figure S5** — Adjusted longitudinal and seasonal trends in sleep timing and duration from mixed-effects models. Model-predicted sleep onset, midsleep, offset, and duration are displayed as a function of longitude. (A) Estimated marginal trends (95% CI) from mixed-effects models, stratified by urbanization status (metropolitan: purple; non-metropolitan: gray) and day type (weekend: solid lines; weekday: dashed lines). (B, C) Average marginal predictions across longitude by solar season for (B) metropolitan and (C) non-metropolitan populations. Slopes reflect fixed-effect estimates for weekend (solid) and weekday (dashed) models, holding all other covariates constant at reference values. (D-F) Corresponding estimates for sleep duration. *Note: To improve seasonal interpretability, sunrise times were manually adjusted to average values at Germany's geographic midpoint on the respective solstice/equinox of each solar season: 06:22 (spring), 05:04 (summer), and 08:19 (winter).*

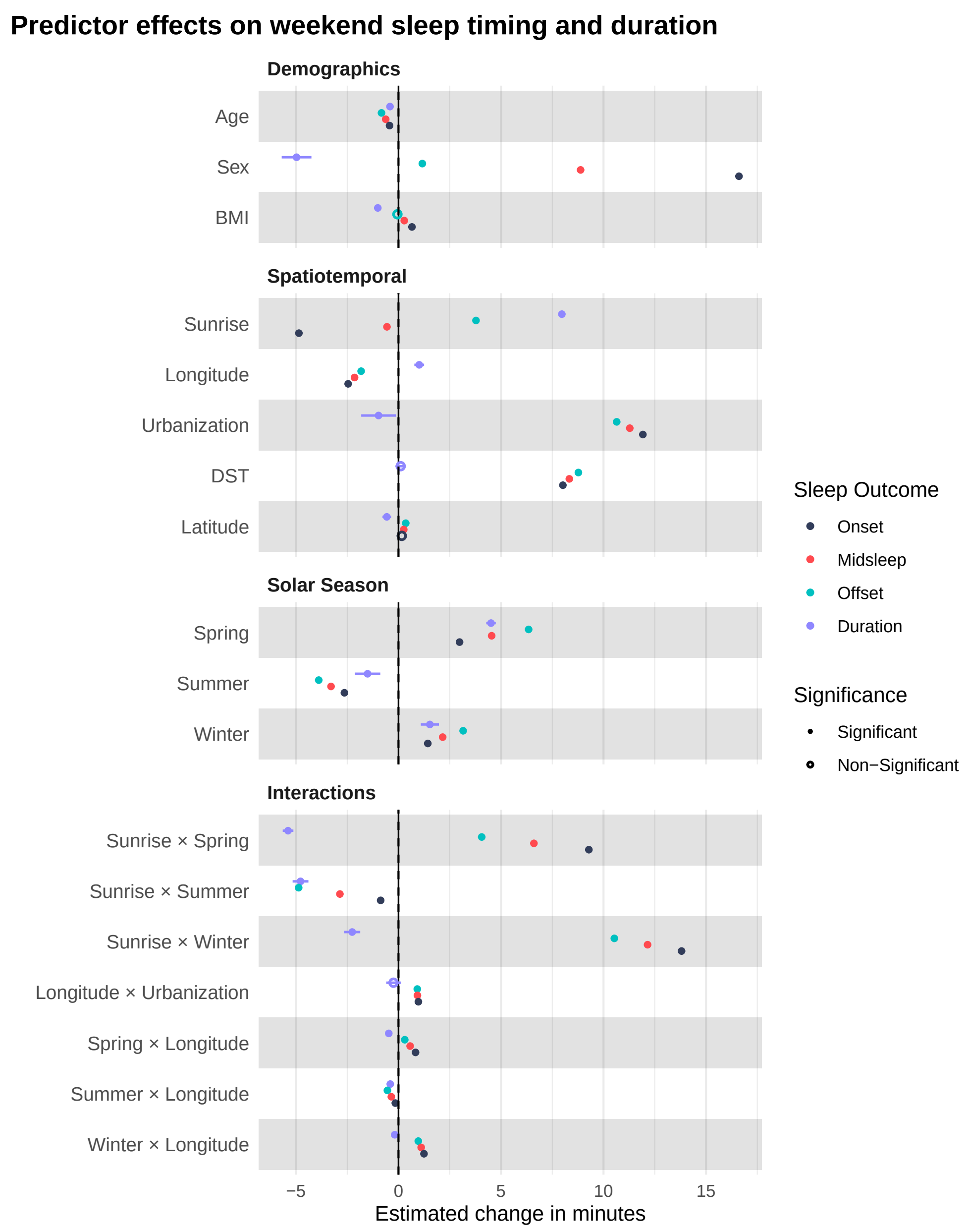


**Figure S6** — Fixed effects from linear mixed-effects models of weekend sleep timing and duration. Forest plot of unstandardized fixed-effect estimates and 95% confidence intervals for predictors of sleep onset, midsleep, offset, and duration on weekends. All models included participant-level random intercepts and were adjusted for demographic, geographic, seasonal, and environmental variables (see Supplementary Methods). Continuous predictors were centered prior to model fitting. Positive estimates indicate later sleep timing or longer sleep duration, while negative estimates indicate earlier sleep or shorter duration. Interaction terms test whether the effects of sunrise and longitude vary by season or urbanization level. Full model results are reported in Table 2.

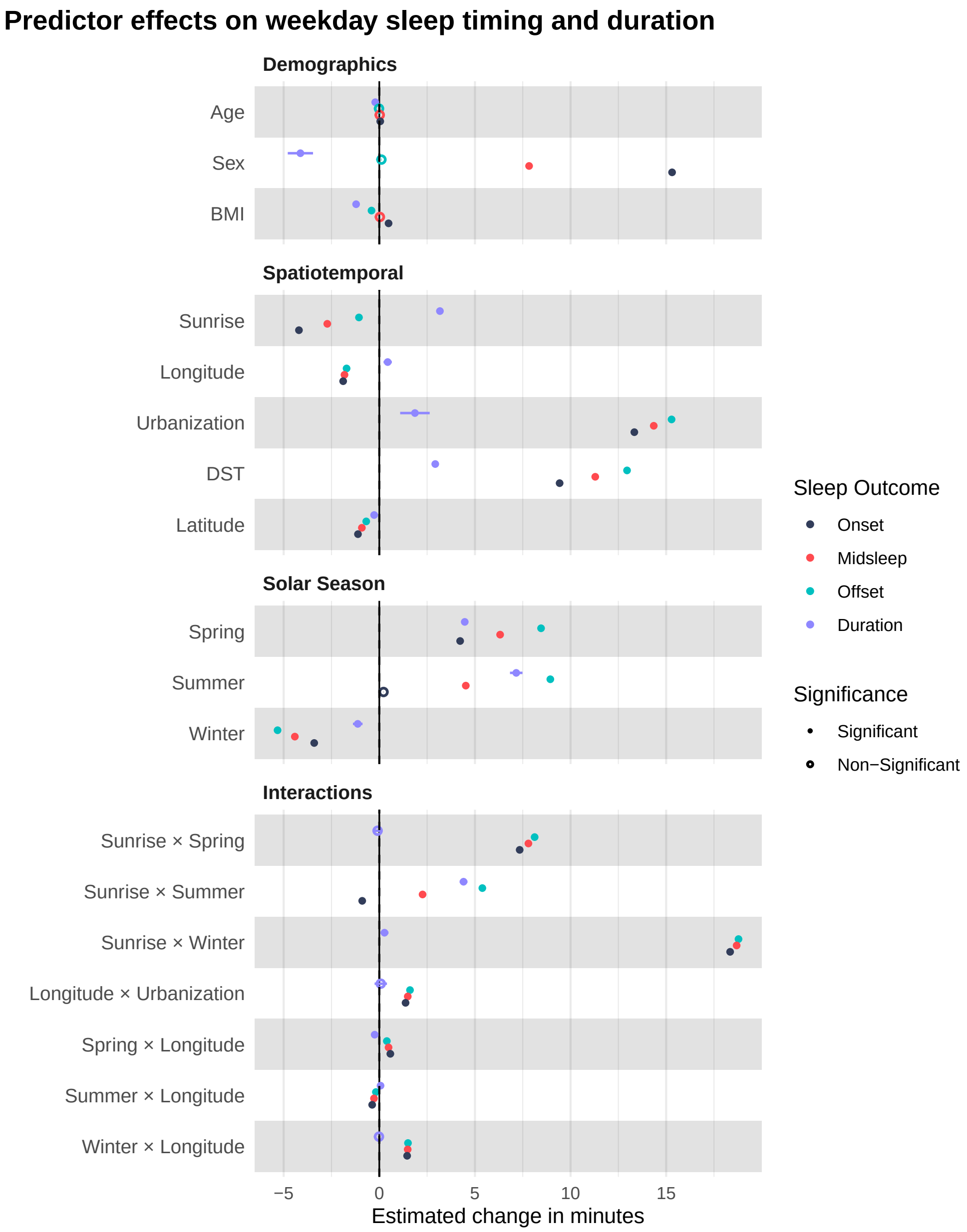


**Figure S7** — Fixed effects from mixed-effects models of sleep timing and duration on weekdays. Forest plot showing unstandardized coefficient estimates and 95% confidence intervals from linear mixed-effects models predicting sleep onset, midsleep, offset, and duration on weekdays. Models included participant-level random intercepts and were adjusted for demographic, geographic, seasonal, and environmental covariates (see Supplementary Methods). Continuous variables were centered prior to model fitting. Positive coefficients indicate later sleep timing or longer sleep duration; negative coefficients indicate earlier timing or shorter duration. Interaction terms assess whether the effects of longitude and sunrise vary by season or urbanization. Model summaries are reported in Supplementary Table 3.

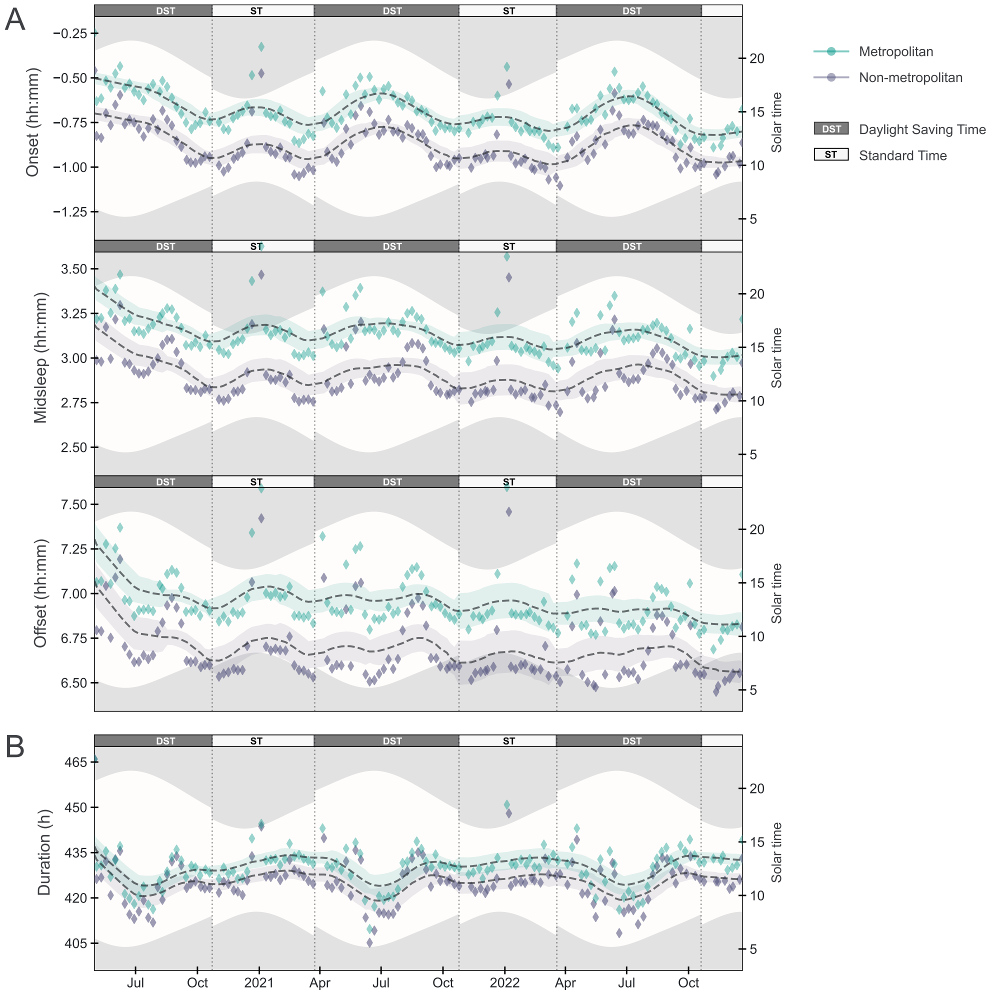


**Figure S8** — Seasonal patterns in weekday sleep timing and duration. (A) Temporal trends in sleep onset, midsleep, offset, and duration over a 20-month period. The smoothed solid lines represent rolling averages of aggregated weekday sleep data. Weekly fluctuations reflect Monday–Friday differences, while broader oscillations align with seasonal changes in solar timing. Vertical dashed lines denote daylight saving time (DST) transitions; shaded bands indicate average photoperiod, bounded by mean sunrise and sunset times. The five days following DST transitions and the winter holiday period (December 24–January 3) were excluded. (B) Annual variation in sleep duration. Duration peaks in winter (standard time; ST) and reaches a minimum in summer (DST), consistent with seasonal changes in both light exposure and social behavior. The amplitude of variation is smaller on weekdays than weekends (see Fig. 3), suggesting that weekday constraints suppress seasonal modulation.

#### Supplementary Tables

**Table S1** — Model comparison for predicting midsleep timing. This table summarizes the stepwise model-building process, beginning with a null model that includes only participant-level random intercepts. The "w/o Demographics" model adds geographic (longitude, latitude, urbanization) and seasonal predictors (season, DST, sunrise time). The "w/ Demographics" model further includes individual-level covariates (age, sex, BMI). The full model incorporates interaction terms to test for seasonal and urbanization-dependent modulation of geographic effects. Model fit is assessed via AIC, BIC, and log-likelihood, with likelihood ratio tests ($\chi^2$) comparing each model to its predecessor. Random effects account for within-participant clustering.

| | **Null Model** | **w/o Demogs** | **w/ Demogs** | **Full Model** |
|---|---|---|---|---|
| Intercept | 229.45 (0.15)*** | 216.35 (0.22)*** | 212.33 (0.35)*** | 209.56 (0.37)*** |
| Sunrise | - | 2.02 (0.03)*** | 2.04 (0.04)*** | -0.56 (0.06)*** |
| Season[spring] | - | 1.72 (0.06)*** | 1.88 (0.07)*** | 4.54 (0.09)*** |
| Season[summer] | - | 6.08 (0.07)*** | 6.05 (0.09)*** | -3.29 (0.24)*** |
| Season[winter] | - | 8.86 (0.07)*** | 8.85 (0.09)*** | 2.15 (0.17)*** |
| DST[yes] | - | 6.34 (0.06)*** | 6.24 (0.08)*** | 8.33 (0.09)*** |
| Longitude | - | -1.36 (0.07)*** | -1.54 (0.10)*** | -2.14 (0.13)*** |
| Latitude | - | 0.08 (0.09) | 0.57 (0.12)*** | 0.26 (0.12)* |
| Urbanization[Urb] | - | 12.17 (0.32)*** | 10.38 (0.43)*** | 11.28 (0.47)*** |
| Age | - | - | -0.62 (0.02)*** | -0.62 (0.02)*** |
| Sex | | | | |
| Sex[male] | - | - | 8.89 (0.40)*** | 8.88 (0.40)*** |
| BMI | - | - | 0.28 (0.04)*** | 0.28 (0.04)*** |
| Spring*Longitude | - | - | - | 0.57 (0.03)*** |
| Summer*Longitude | - | - | - | -0.35 (0.03)*** |
| Winter*Longitude | - | - | - | 1.11 (0.03)*** |
| Longitude*Urban | - | - | - | 0.92 (0.20)*** |
| Spring*Sunrise | - | - | - | 6.60 (0.10)*** |
| Summer*Sunrise | - | - | - | -2.86 (0.15)*** |
| Winter*Sunrise | - | - | - | 12.15 (0.15)*** |
| *Observations* | 12,876,804 | 12,876,804 | 7,315,838 | 7,315,838 |
| *Num. Users* | 105,729 | 105,729 | 59,767 | 59,767 |
| *Log-Likelihood* | -16,935,389 | -16,914,327 | -9,550,357 | -9,544,949 |
| *Akaike Inf. Crit.* | 33,870,784 | 33,828,675 | 19,100,742 | 18,909,948 |
| *Bayesian Inf. Crit.* | 33,870,827 | 33,828,833 | 19,100,935 | 18,909,293 |
| *Marg./Cond.* $R^2$ | 0.00/0.46 | 0.01/0.45 | 0.02/0.47 | 0.02/0.47 |

*Note:* $^{*}$p<0.05; $^{**}$p<0.01; $^{***}$p<0.001

**Table S2** — Sensitivity analyses for mixed models predicting midsleep. The table evaluates robustness to alternative model specifications, comparing the base model to variations addressing potential biases. The "Random Effects (Day)" model includes an additional random effect for day-level variation. The "Alt Sunrise Centering" model tests the impact of shifting the sunrise reference point. The "Dual Sunrise Var" model incorporates both absolute and relative sunrise times. The "East-West Var" model accounts for regional effects by including an additional longitudinal variability term. Model fit is assessed using AIC, BIC, and likelihood ratio tests ($\chi^2$) comparing each model against the base model. Lower AIC and BIC values indicate better model fit. Fixed effects are reported as estimates with standard errors in parentheses.

| | **Base Model** | **Random Effects (Day)** | **Alt Sunrise Centering** | **Dual Sunrise Vars** | **East-West Var** |
|---|---|---|---|---|---|
| Intercept | 212.3 (0.35)*** | 211.3 (2.40)*** | 212.6 (0.35)*** | 210.9 (0.43)*** | 212.6 (0.77)*** |
| Sunrise | 2.04 (0.04)*** | 2.70 (0.20)*** | 2.04 (0.04)*** | -0.44 (0.44)*** | 2.04 (0.04)*** |
| Season[spring] | 1.88 (0.07)*** | 1.89 (0.07)*** | 0.40 (0.07)*** | 1.88 (0.07)*** | 1.88 (0.07)*** |
| Season[summer] | 6.05 (0.09)*** | 7.11 (0.09)*** | 1.93 (0.06)*** | 6.05 (0.09)*** | 6.05 (0.09)*** |
| Season[winter] | 8.85 (0.09)*** | 8.30 (0.10)*** | 11.32 (0.10)*** | 8.85 (0.09)*** | 8.85 (0.09)*** |
| DST[yes] | 6.24 (0.08)*** | 6.52 (0.08)*** | 6.24 (0.08)*** | 6.24 (0.08)*** | 6.24 (0.08)*** |
| Longitude | -1.54 (0.10)*** | -1.49 (0.10)*** | -1.54 (0.10)*** | -1.70 (0.10)*** | -1.57 (0.12)*** |
| Latitude | 0.57 (0.12)*** | 0.56 (0.12)** | 0.57 (0.12)** | 0.51 (0.12)** | 0.56 (0.12)** |
| Urbanization[urb] | 10.38 (0.43)*** | 10.44 (0.43)*** | 10.38 (0.43)*** | 10.41 (0.43)*** | 10.36 (0.43)*** |
| Age | -0.62 (0.02)*** | -0.63 (0.02)*** | -0.62 (0.02)*** | -0.63 (0.02)*** | -0.62 (0.02)*** |
| Sex[male] | 8.89 (0.40)*** | 9.15 (0.40)*** | 8.89 (0.40)*** | 9.06 (0.40)*** | 8.89 (0.40)*** |
| BMI | 0.28 (0.04)*** | 0.28 (0.04)*** | 0.28 (0.04)*** | 0.28 (0.04)*** | 0.28 (0.04)*** |
| *Observations* | 7,315,838 | 7,315,838 | 7,315,838 | 7,315,838 | 7,315,838 |
| *Num. Users* | 59,767 | 59,767 | 59,767 | 59,767 | 59,767 |
| *Log-Likelihood* | -9,544,949 | -9,456,000 | -9,544,949 | -9,550,341 | -9,550,263 |
| *Akaike Inf. Crit.* | 18,909,948 | 18,912,053 | 18,909,948 | 18,907,112 | 18,900,562 |
| *Bayesian Inf. Crit.* | 18,909,293 | 18,912,412 | 18,909,293 | 18,909,919 | 18,900,811 |
| *Marg./Cond.* $R^2$ | 0.02/0.47 | 0.02/0.47 | 0.02/0.47 | 0.02/0.47 | 0.02/0.47 |
| *Note:* | | | | | $^{*}$p<0.05; $^{**}$p<0.01; $^{***}$p<0.001 |

**Table S4** — Linear mixed model results for sleep timing and duration on weekdays. Each column reports unstandardized fixed-effect estimates (with 95% confidence intervals) from models predicting sleep onset, midsleep, offset, and duration. Models include participant-level random intercepts and adjust for demographic (age, sex, BMI), geographic (longitude, latitude, urbanization), seasonal (season, DST), and environmental (sunrise time) variables, as well as interactions between geography and season. Positive coefficients indicate later sleep timing or longer duration; negative coefficients indicate earlier timing or shorter duration. $R^2$ values denote variance explained by fixed effects alone (marginal) and by the full model including random effects (conditional). Statistical significance is denoted by asterisks.

| | *Sleep Variable* | | | |
|---|---|---|---|---|
| | ONSET | MIDSLEEP | OFFSET | DURATION |
| Intercept | $-75.780^{***}$ | $155.580^{***}$ | $387.480^{***}$ | $421.847^{***}$ |
| | (-76.49, -75.07) | (154.87, 156.29) | (386.77, 388.19) | (421.24, 422.45) |
| Age | $0.060^{**}$ | 0.000 | 0.000 | $-0.212^{***}$ |
| | ( 0.02, 0.10) | ( 0.00, 0.00) | ( 0.00, 0.00) | (-0.24, -0.19) |
| Sex[male] | $15.300^{***}$ | $7.860^{***}$ | 0.120 | $-4.125^{***}$ |
| | (14.48, 16.12) | ( 7.15, 8.57) | (-0.70, 0.94) | (-4.79, -3.46) |
| BMI | $0.480^{***}$ | 0.060 | $-0.420^{***}$ | $-1.212^{***}$ |
| | ( 0.36, 0.60) | (-0.06, 0.18) | (-0.54, -0.30) | (-1.28, -1.15) |
| Urbanization[urban] | $13.320^{***}$ | $14.340^{***}$ | $15.300^{***}$ | $1.863^{***}$ |
| | (12.38, 14.26) | (13.22, 15.46) | (14.40, 16.20) | ( 1.09, 2.64) |
| DST[yes] | $9.420^{***}$ | $11.280^{***}$ | $12.960^{***}$ | $2.926^{***}$ |
| | ( 9.30, 9.54) | (11.16, 11.40) | (12.84, 13.08) | ( 2.78, 3.07) |
| Season[spring] | $4.200^{***}$ | $6.300^{***}$ | $8.460^{***}$ | $4.468^{***}$ |
| | ( 4.08, 4.32) | ( 6.18, 6.42) | ( 8.34, 8.58) | ( 4.32, 4.61) |
| Season[summer] | 0.240 | $4.500^{***}$ | $8.940^{***}$ | $7.160^{***}$ |
| | (-0.24, 0.48) | ( 4.27, 4.73) | ( 8.59, 9.29) | ( 6.83, 7.49) |
| Season[winter] | $-3.420^{***}$ | $-4.440^{***}$ | $-5.340^{***}$ | $-1.126^{**}$ |
| | (-3.66, -3.18) | (-4.68, -4.20) | (-5.58, -5.10) | (-1.38, -0.88) |
| Longitude | $-1.860^{***}$ | $-1.800^{***}$ | $-1.740^{***}$ | $0.439^{***}$ |
| | (-2.10, -1.62) | (-2.04, -1.56) | (-1.98, -1.50) | ( 0.22, 0.66) |
| Latitude | $-1.140^{***}$ | $-0.900^{***}$ | $-0.660^{***}$ | $-0.269^{**}$ |
| | (-1.38, -0.90) | (-1.14, -0.66) | (-0.90, -0.42) | (-0.46, -0.08) |
| Sunrise | $-4.200^{***}$ | $-2.700^{***}$ | $-1.080^{***}$ | $3.170^{***}$ |
| | (-4.32, -4.08) | (-2.82, -2.58) | (-1.20, -0.96) | ( 3.09, 3.25) |
| Spring:Lon | $0.600^{***}$ | $0.480^{***}$ | $0.420^{***}$ | $-0.241^{***}$ |
| | ( 0.60, 0.60) | ( 0.48, 0.48) | ( 0.42, 0.42) | (-0.28, -0.20) |
| Summer:Lon | $-0.360^{***}$ | $-0.300^{***}$ | $-0.180^{***}$ | $0.066^{**}$ |
| | (-0.36, -0.36) | (-0.30, -0.30) | (-0.18, -0.18) | ( 0.03, 0.10) |
| Winter:Lon | $1.440^{***}$ | $1.500^{***}$ | $1.500^{***}$ | $-0.017$ |
| | ( 1.44, 1.44) | ( 1.50, 1.50) | ( 1.50, 1.50) | (-0.06, 0.02) |
| Lon:Urban | $1.380^{***}$ | $1.500^{***}$ | $1.620^{***}$ | 0.077 |
| | ( 1.03, 1.73) | ( 1.15, 1.85) | ( 1.27, 1.97) | (-0.25, 0.40) |
| Spring:Sunrise | $7.320^{***}$ | $7.800^{***}$ | $8.100^{***}$ | $-0.089$ |
| | ( 7.20, 7.44) | ( 7.68, 7.92) | ( 7.98, 8.22) | (-0.24, 0.06) |
| Summer:Sunrise | $-0.900^{***}$ | $2.280^{***}$ | $5.400^{***}$ | $4.405^{***}$ |
| | (-1.14, -0.66) | ( 2.16, 2.40) | ( 5.17, 5.63) | ( 4.20, 4.61) |
| Winter:Sunrise | $18.360^{***}$ | $18.660^{***}$ | $18.780^{***}$ | $0.272^{**}$ |
| | (18.12, 18.60) | (18.54, 18.78) | (18.55, 19.02) | ( 0.06, 0.48) |
| *Observations* | 18,299,364 | 18,403,489 | 18,399,909 | 18,500,002 |
| *Num. Users* | 59,772 | 59,772 | 59,772 | 59,772 |
| *Log Likelihood* | -11,179,576 | -9,636,392 | -10,344,812 | -41,682,215 |
| *AIC* | 22,359,207 | 19,272,881 | 20,689,706 | 83,368,444 |
| *BIC* | 22,359,497 | 19,273,171 | 20,689,996 | 83,368,734 |
| *Marg./Cond.* $R^2$ | 0.03/0.48 | 0.02/0.50 | 0.01/0.45 | 0.01/0.33 |

*Note:* $^{*}p < 0.05$; $^{**}p < 0.01$; $^{***}p < 0.001$
